# POTTR: Identifying Recurrent Trajectories in Evolutionary and Developmental Processes using Posets

**DOI:** 10.64898/2026.02.25.707960

**Authors:** Sara C. Käufler, Henri Schmidt, Martin Jürgens, Gunnar W. Klau, Palash Sashittal, Benjamin J. Raphael

## Abstract

Multiple biological processes, including cancer evolution and organismal development, are described as a sequence of events with a temporal ordering. While cancer evolves independently in each patient, DNA sequencing studies have shown that in some cancers different patients share specific orders of mutations and these correlate with distinct morphology, drug response, and treatment outcomes. Several methods have been developed to identify such recurrent trajectories of genetic events from phylogenetic trees, but this is complicated by high intra- and inter-tumor heterogeneity as well as uncertainty in the inferred tumor phylogenies including the ambiguous orders between some mutations. We formalize the problem of finding recurrent mutation trajectories using a novel framework of *incomplete* partially ordered sets (posets), which generalize representations used in previous works and explicitly account for the uncertainty in tumor phylogenies. We define the problem of identifying the largest recurrent trajectories shared in at least *k* input phylogenies as the maximum *k*-common induced incomplete subposet (MkCIIS) problem, which we show is NP-hard. We present a combinatorial algorithm, POsets for Temporal Trajectory Resolution (POTTR), to solve the MkCIIS problem using a conflict graph that models recurrent trajectories as independent sets. Thereby we identify maximum recurrent trajectories while resolving multiple sources of uncertainty, like mutation clusters, in the phylogenetic data. We apply POTTR to TRACERx non-small cell lung cancer bulk sequencing and acute myeloid leukemia single-cell sequencing data and through resolution of mutation clusters discover previously unreported trajectories of high statistical significance. On lineage tracing data of an in vitro embryoid model, POTTR identifies conserved differentiation routes across biological replicates and how these routes change in response to chemical perturbations.

## 1 Introduction

Many biological processes involve a progression of heritable events in cells over time. For example, cells accumulate somatic mutations as they divide, sometimes leading to cancer, or differentiate into new cell types during organismal development. As it is often not possible to measure cells longitudinally from the same organism, processes such as somatic evolution and development are studied by using phylogenetic techniques to infer past events based on the measurement of a population of cells at a single time. For example, many methods have been developed to infer rooted phylogenetic trees that describe cancer evolution [13,14,15,16,31,46,54,60,61] or that infer cell lineage trees [34,18,22,55,11]. In these trees, nodes commonly denote mutations, subclones, or cell types, and edges represent their sequential occurrence over time. From such a tree, one can infer ancestral events and their order in *one* individual. But due to stochasticity in evolutionary processes (and some developmental processes [50]) an important question is to derive recurrent trajectories, which are orders of events that are shared across individuals.

The problem of finding recurrent trajectories across individuals is arguably best studied for the somatic evolutionary process in cancer. Tumor formation is a complex evolutionary process whereby the tumor is composed of different subpopulations, *clones*, of cells that differ in their mutation profiles [49]. During its growth, the tumor undergoes a series of selection and expansion steps, which result in high intratumoral as well as high intertumoral heterogeneity, presenting one of the biggest challenges in cancer treatment. Despite this considerable diversity, through advances in sequencing and phylogeny inference methods it has been shown that certain tumors follow common evolutionary patterns across patients. Although shared evolutionary trajectories can be observed, most cancer driver genes are mutated only in a small fraction of patients with a given cancer type therefore limiting the size of the trajectories [58]. Multiple studies provided evidence that the order in which the mutations were accumulated by the tumor play a crucial role in disease and treatment outcome [51,36,41,20]. Ortmann et al. [51], for example, examined the temporal order effect of Janus kinase 2 (*JAK2*) and *TET2* mutations in myeloproliferative neoplasm patients. Both mutations were present in about 10% of their cohort. Patients that accumulated the *TET2* mutation first, were on average older and their neoplasms were smaller in size compared to patients in which *JAK2* mutated first. Moreover, in vitro colony formation experiments revealed differences in how tumor cells respond towards treatment with a *JAK2* inhibitor. The cells, in which the *JAK2* mutation occurred first, were much more sensitive to the inhibitor than cells, in which the *TET2* mutation was present first [51].

The problem of finding recurrent trajectories depends on the definition of a trajectory and the properties of the input data. A first important issue is whether the recurrent trajectories are linear orders or branching trajectories. A second issue is whether all events in each trajectory are distinguishable or if some events are grouped into indistinguishable clusters. For cancer phylogenies, we define a recurrent trajectory as a set of mutations on which the same partial order is observed in phylogenetic trees from distinct tumors (Fig. 1). However, due to technical limitations in sequencing depth or sampling of cells, some mutations may appear together in the same set of cells/clones within a patient. Importantly, the mutations on a recurrent trajectory need to be consecutive in each patient, but might have intervening events in some patients that are not part of the recurrent trajectory. In addition, due to uncertainty in tree inference, there may be multiple plausible trees for a patient, which methods should take into account to ensure robust results.

**Fig. 1.**
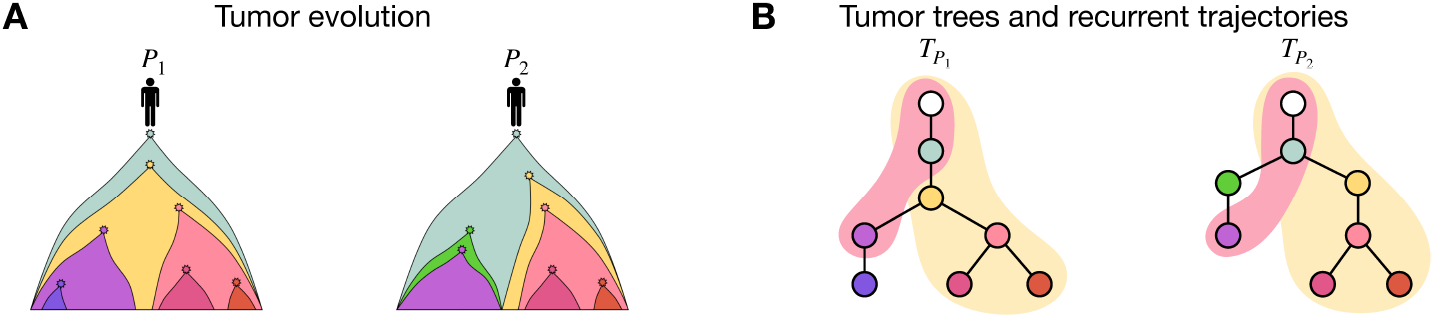
A Illustration of tumor evolution in two patients *P*_1_ and *P*_2_, colors mark mutation events. In patient *P*_1_ the blue and yellow mutation are clonal, while in *P*_2_ the blue mutation is clonal and the yellow mutation is subclonal. **B** The corresponding tumor trees. The white node is the root node, indicating the normal cells. Highlighted in pink and yellow are two distinct recurrent trajectories, in which the purple and yellow node have different orderings.

Multiple computational methods to identify recurrent trajectories from cancer phylogenetic trees have been developed which we classify into two categories (Table 1). The first, feature-based methods, learn statistical or probabilistic models over the entire set of mutations to identify common evolutionary patterns. REVOLVER [8] and Hintra [37] both resolve clusters and consider multiple phylogenies per patient, but REVOLVER analyzes only recurrent linear orders while Hintra extends the approach to branching structures. However, since Hintra enumerates all possible tree topologies it does not scale well to large input data, i.e. trees with more than five alterations [52]. TreeMHN [44] adapts the idea of jointly inferring recurrent trajectories, but takes as input a single tree per patient and is restricted to linear clonal structures instead of capturing branching evolutionary trajectories. Moreover, REVOLVER, Hintra, and TreeMHN are limited by their use of a single model to describe all phylogenies simultaneously and do not account for distinct subtypes of tumors, which are known to evolve differently [52,10]. GeneAccord [39] is a statistical method that focuses on pairwise relations and mutual exclusivity that are statistically significant without drawing any conclusions from branching substructures. Ivanociv et al. [30] present a tree generative model, CloMu, that probabilistically infers recurrent trajectories of interchangeable mutations from multiple input trees per patient. Oncotree2vec [3] clusters trees with similar evolutionary patterns, however, which substructures are responsible for the cluster is generally open to interpretation.

**Table 1.** Properties of recurrent trajectory inference that are modeled by different methods.

|  | Feature-based methods |  |  | Alignment-based methods |  |  |  |
| --- | --- | --- | --- | --- | --- | --- | --- |
|  | REVOLVER | Hintra | CloMu | RECAP | CONETT | MASTRO | POTTR |
| Branching trajectories |  | ✓ | ✓ | ✓ | ✓ | ✓ | ✓ |
| Resolving clusters | ✓ | ✓ | ✓ | ✓ | ✓ |  | ✓ |
| Multiple trees per patient | ✓ | ✓ | ✓ | ✓ |  |  | ✓ |
| Induced subgraphs |  |  |  |  |  | ✓ | ✓ |
| Exact |  |  |  |  | ✓ | ✓ | ✓ |

The second category of methods are tree alignment methods, which explicitly compare and align trees to report a common consensus or conserved orders of mutations. Consensus tree methods [23,1] summarize the input trees by minimizing a defined tree distance and thereby capture common structures, which makes them useful for clustering tasks. However, while consensus trees are similar to real trees, they are a mosaic of multiple trees and thus do not necessarily describe real tumor evolution or conserved orders over mutations. RECAP [10] and CONETT [28] resolve mutation clusters and compute common consensus trees. RECAP-considers multiple trees per tumor and heuristically selects and clusters tumor phylogenetic trees, and infers a consensus tree per cluster. In contrast, CONETT considers one tree per tumor and uses an integer linear program (ILP) to infer a single consensus tree for the largest possible set of input trees. Instead of computing consensus trees, Pellegrina and Vandin [52] propose a method, MASTRO, which models recurrent trajectories as common induced subgraphs of tumor trees, another important property of methods for identifying recurrent trajectories. MASTRO uses frequent itemset mining to find all maximal recurrent trajectories. However, MASTRO cannot deal with multiple input trees per patient and does not resolve mutation clusters. Finally, some of the above methods solve their formulation of the problem of finding recurrent trajectories exactly, while others rely on approximations or heuristic approaches.

We recast the problem of finding recurrent trajectories in terms of partially ordered sets (posets), which can be used to present temporal processes, such as tumor evolutionary histories, but also to model cellular development, where they define the order of differentiation events. This formulation generalizes many of the existing works on identifying recurrent trajectories and has two major advantages. First, this extends the problem beyond trees to partial orders that are not tree-like, but rather their transitive reduction is a directed acyclic graph (DAG). Second, the formulation enables a rigorous description of the problem of resolving events with indistinguishable order (i.e. clusters of mutation) using a novel structure that we call an incomplete poset. We propose the *Maximum k-Common Induced Incomplete Subposet (MkCIIS) Problem* to identify the largest set of events that follow the same (incomplete) partial order in at least *k* input posets. The user-defined parameter *k* allows for identifying orders of events that are conserved in only a fraction of the input. We show that MkCIIS is NP-hard and reformulate it as the problem of finding an independent set in a multi-edge-conflict-graph. We derive a combinatorial method, POTTR, to solves MkCIIS using an integer linear program (ILP). We applied POTTR to 401 non-small cell lung cancer (NSCLC) patients from TRACERx [19] and found a number of statistically significant recurrent trajectories complying with previously reported results and extending these through resolving mutation clusters. We also compared POTTR to MASTRO on an earlier TRACERx data set of 99 NSCLC trees and 123 trees of acute myeloid leukemia (AML) single-cell panel sequencing data provided by Pellegrina and Vandin [52], which were originally obtained from [32] and [47], respectively. We show that POTTR can handle multiple sources of uncertainty and by resolving these uncertainties, discover previously unreported trajectories of high biological relevance. Lastly, we applied POTTR on lineage tracing data of Trunk-Like Structures (TLS), an in vitro embryoid model, to identify conserved differentiation routes across biological replicates and how these routes change in response to chemical perturbations.

## 2 Recurrent Trajectories and Incomplete Posets

For the systematic comparison of tumor evolution across patients, the standard approach is to develop algorithms that operate on the rooted phylogenetic tree(s) of each patient. Rooted phylogenetic trees describe a *partial order* on the mutations within a tumor, where a mutation *a* precedes a mutation *b* (written as *a* ≺ *b*) if *a* is on the unique path from *b* to the root node. However, trees are only a special case of a partial order (more generally partial orders correspond to DAGs) and it is not always clear how operations on trees affect the underlying partial order.

Consequently, we describe the problem of identifying recurrent trajectories using the framework of partial orders and partially ordered sets (posets). Given a set *S* of elements, a *partial order* ≺ is a relation that is reflexive, asymmetric, and transitive, while a *strict* partial order is irreflexive, asymmetric and transitive. In this paper, we will use partial order to indicate a strict partial order. The tuple *P* = (*S*, ≺) is also referred to as partially ordered set, or *poset*. In contrast to total orders, partial orders do not require all elements to be ordered but allow elements to be incomparable.

An additional complication arises in DNA sequencing data of tumors in that the (partial) order of individual mutations is not always known in a single patient. Instead, groups of mutations are measured together in the same clones or cells. In tumor phylogenetic trees, mutations whose relative order cannot be determined, are typically grouped into a single node called a mutation cluster, which implicitly models an underlying linear order on these mutations. We address this additional uncertainty in the language of partially ordered sets by introducing the concept of indistinguishable elements, or *hidden orders* ∼, on a set of elements *S*. The hidden order is an equivalence relation, which is reflexive, symmetric, and transitive, such that if *a* ∼ *b* and *b* ∼ *c*, then *a* ∼ *c*. Like any equivalence relation, ∼ defines a partition *P*_H_ of *S* into hidden sets, where each hidden set consists of elements related by the hidden order.

For a set *S*, we define an *incomplete partial order* on *S* as the pair (∼, ≺), where ∼ is a hidden order on *S* and ≺ is a partial order on the elements of the partition *P*_ℋ_ induced by ∼ on *S*. The sets of the partition *P*_ℋ_ are related by a strict partial order, which passes on to elements from distinct sets of the partition. This is, for any pair *H*_1_, *H*_2_ ∈ *P*_ℋ_ with *H*_1_ ≺ *H*_2_, *a* ≺ *b* for every *a* ∈ *H*_1_, *b* ∈ *H*_2_. We call *P* = (*S*, ∼, ≺) an *incomplete poset*. Incomplete partial orders can be thought of as an extension of strict partial orders that allow the ordering of some subsets of elements to be unknown/unresolved. Although partial orders on equivalence relations have been studied [2,29], to the best of our knowledge, incomplete partial orders, where each hidden order has an underlying linear order, are an unexplored concept. We can resolve the order of indistinguishable elements of a hidden set by splitting the set into two ordered subsets. We define a resolution of an incomplete poset as follows.

### Definition 1.

*Let P* = (*S*, ∼, ≺) *be an incomplete poset on S and let H* ∈ *P*_ℋ_ *be a hidden set. P*′ = (*S*, ∼′, ≺′) *is a* ***resolution*** *of P if there is a partition H* = *J*_1_∪*J*_2_ *of H with J*_1_, *J*_2_ ≠ ∅ *such that:* 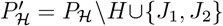; *J*_1_ ≺′ *J*_2_; *all sets G* ∈ *P*_ℋ_ *preceding H* ∈ *P precede both J*_1_ *and J*_2_ *in P*′, *and all sets G* ∈ *P*_ℋ_ *succeeding H* ∈ *P succeed both J*_1_ *and J*_2_ *in P*′.

The resolution of an incomplete poset is an incomplete poset as well. If we resolve all hidden orders of an incomplete poset, that is, 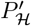 consists of singletons only, we would get a poset. We refer to this as a *completion* of an incomplete poset.

Given a family of incomplete posets as input, we search for recurrent evolutionary trajectories. More precisely, we seek to find a subset of events with an (incomplete) partial order such that the following criteria are met:

- For at least *k* incomplete posets in the family, all orders and incomparabilities on this subset are preserved, while hidden orders can be resolved.
- The selected subset of events is of maximum size, i.e., no larger subset of events exists for which the order is preserved across *k* incomplete posets.

To identify recurrent trajectories it is important to consider the transitivity of incomplete posets. Specifically, not all tumor trees from a patient cohort share the same set of mutations. Transitivity allows us to skip over mutations that are not shared and enables finding recurrent trajectories despite differences across tumor trees.

To identify shared ordered events, we look at the intersection of multiple incomplete posets. Given a family of incomplete posets 𝒫 = {(*S*_1_, ∼ _1_, ≺ _1_),…, (*S*_*n*_, ∼ _*n*_, ≺ _*n*_)}, the *intersection* (*S*, ∼, ≺) over 𝒫 is the incomplete poset where 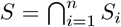, and for all *a, b* ∈ *S, a* ≺ *b* if and only if *a* ≺ _*i*_ *b* and *a* ∼ *b* if and only if *a* ∼ _*i*_ *b* for all *i* = 1,…, *n*. The intersection yields a unique set of common elements and pairwise (hidden) orders. However, the intersection might ignore certain relations that are relevant for an evolutionary process.

We describe this issue in Fig. 2 using an example and the graph representation of posets, which are equivalent to transitively closed DAGs [57]. Posets are often represented in form of a Hasse diagram, which is the transitive reduction of that DAG. Incomplete partial orders also have an equivalent representation as transitively closed DAGs, in which each set of the partition *P*_ℋ_ maps to a node and precedence relations between sets define the directed edges between the corresponding nodes. The resolution of a cluster also has a clear interpretation in the graph representation: when a cluster is resolved, the node representing that cluster is split into two nodes that are connected by a directed edge, reflecting the outcome of the resolution. In the following figures, for the sake of visualization, we will represent posets by their equivalent graph representations. As depicted in Fig. 2 B, the orders “green precedes yellow” and “green precedes pink” are shared in all three input trees. The contradictory ordering of yellow and pink in the input graphs in panel A is not reflected here. This shows that the intersection is not necessarily a recurrent order.

**Fig. 2.**
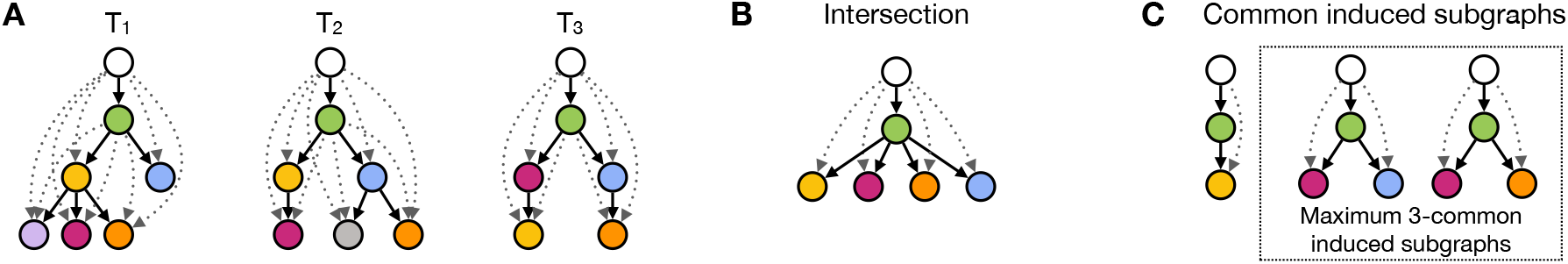
Comparison of graph intersection and common induced subgraphs. **A** Three trees *T*_1_, *T*_2_, *T*_3_ with tree edges (solid lines) and the transitive partial order relations (dotted arrows). **B** The unique intersection of *T*_1_, *T*_2_, and *T*_3_. **C** Three common induced subgraphs of *T*_1_, *T*_2_, and *T*_3_. The graphs in the box are maximum 3-common induced subgraphs based on their number of nodes.

For a single incomplete poset, we interpret a recurrent order on a subset of the elements as an *induced incomplete subposet* and define it analogously to the induced subposets (or sometimes restrictions) described by Bukh [7]. Similarly, we can define *common induced incomplete subposets* for a family𝒫 of incomplete posets.

### Definition 2.

*P* = (*S*, ∼, ≺) *is an* ***induced incomplete subposet*** *of an incomplete poset P*′ = (*S*′, ∼′, ≺′) *provided S* ⊆ *S*′ *and for all a, b* ∈ *S, a* ≺ *b if and only if a* ≺′ *b, and a* ∼ *b if and only if a* ∼′ *b*.

### Definition 3.

*P is a* ***common induced incomplete subposet (CIIS)*** *of a family* 𝒫 *of incomplete posets if P is an induced incomplete subposet of each incomplete poset in* 𝒫.

In contrast to the intersection, there could be multiple CIIS with varying numbers of elements, see Fig. 2 C. For example, each single element that is shared in all the input posets is a CIIS. Moreover, all CIIS of a family of incomplete posets 𝒫 are subposets of the intersection of 𝒫. As such, the size of the largest CIIS of a family of incomplete posets is bounded from above by 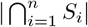.

Following the criteria above and considering the ambiguity of the data, we focus on finding a maximum common induced incomplete subposet among *k* selected incomplete posets from the family 𝒫, where *k* is a user-defined integer. When identifying recurrent trajectories, we can use the information of pairwise orders across incomplete posets to resolve hidden orders among the selected posets. Thus, for each selected poset, the common induced subposet is then an induced subposet of either the poset itself or one of its resolutions. We formally pose the problem as follows and show that it is NP-hard. See Suppl. Mat. Sect. S2.1 for a proof.

### Problem 1

**(Maximum** *k***-Common Induced Incomplete Subposet Problem, (MkCIIS))** *Given a family* 𝒫 = {*P*_1_,…, *P*_*n*_} *of incomplete posets, and an integer k, find a largest poset P and a selection of at least k posets from* 𝒫, *such that P is an induced subposet of each selected poset itself or one of its resolutions*.

### Theorem 1.

*The decision variant of the MkCIIS problem is NP-complete for* 2 ≤ *k* ≤ *n*.

## 3 POTTR: Combinatorial Method for Solving MkCIIS using a Conflict Graph

We introduce our method POsets for Temporal Trajectory Resolution (POTTR), which solves the MkCIIS problem and addresses multiple sources of uncertainty in the data including resolution of hidden orders and the selection among multiple incomplete posets obtained from tumor phylogenetic inference. We use all trees reported for a tumor by phylogenetic inference methods and only decide on a single phylogeny the moment we find a recurrent trajectory. Although, POTTR can take multiple incomplete posets for each evolutionary process as input, for simplicity, we describe our algorithm on input data consisting of one incomplete poset per evolutionary process and defer the full description to Suppl. Mat. Sect. S1.

We derive a combinatorial characterization of common induced incomplete subposets (CIIS) of a family 𝒫 of incomplete posets by introducing the *conflict graph C*(𝒫). The conflict graph *C*(𝒫) is a multigraph that encodes if there are conflicts in the order of events described by any pair of incomplete posets in 𝒫. We demonstrate a one-to-one bijection between CIIS of 𝒫 and independent sets in the conflict graph *C*(𝒫), providing a complete characterization of the combinatorial structure of CIIS.

A conflict exists between two elements of distinct incomplete posets if the relation of these elements is different in each incomplete poset. We formally define the conflict graph *C*(𝒫) for 𝒫 as follows.

### Definition 4

**(Conflict Graph)**. *A conflict graph C*(𝒫) = (*V, E*) *for a family* 𝒫 = {*P*_1_,…, *P*_*n*_} *of incomplete posets is an undirected multigraph whose vertex set* 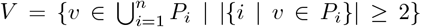 *consists of all elements that occur in at least two distinct incomplete posets. There is an edge* {*a, b*} *in E with label* {*i, j*} *if for a pair P*_*i*_ = (*S*_*i*_, ∼ _*i*_, ≺ _*i*_) *and P*_*j*_ = (*S*_*j*_, ∼ _*j*_, ≺ _*j*_) *of incomplete posets from* 𝒫 *and a pair a, b* ∈ *P*_*i*_, *P*_*j*_, *at least one of the following conflicts exists: (i) a* ≺ _*i*_ *b and b* ≺ _*j*_ *a; or (ii) a* ≺ _*i*_ *b and a and b are incomparable in P*_*j*_; *or (iii) a* ∼ _*i*_ *b and a and b are incomparable in P*_*j*_.

Figure 3 shows four input incomplete posets (A), the corresponding conflict graph (B), and a maximum common induced subposet (Fig. 3C) for three selected input posets. The corresponding nodes form an independent set in the conflict graph for the selected incomplete posets. This is, the nodes are not connected by any edge labeled by any pair of the the selected incomplete posets. Since the conflict graph *C*(𝒫) is constructed on all pairwise conflicts between incomplete posets of a family 𝒫, the conflict graph of a subfamily 𝒫 ⊆ 𝒫 is a subgraph of *C*(𝒫). We derive a bijection between a CIIS and independent sets in the conflict graph in the following proposition (proof in Suppl. Mat. Sect. S2.2).

**Fig. 3.**
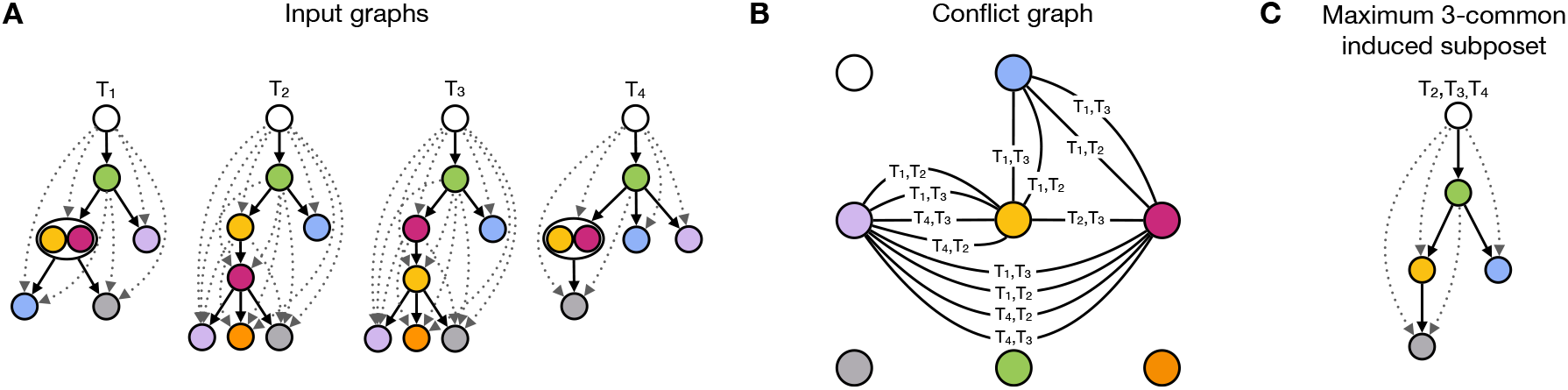
Example of the conflict graph for three sets of input posets. **A** Three distinct sets of input posets presented as transitively closed DAGs. Set 1 consists of two posets, which both have a hidden order for the yellow and pink element. The hidden orders are represented as clusters in the DAGs. **B** Conflict graph with all pairwise conflicts between posets of distinct sets. **C** A maximum common induced subposet for the selection of *T*_2_, *T*_3_, and *T*_4_.

### Proposition 1.

*Let 𝒫 be a family of incomplete posets and let C*(𝒫) *be the corresponding conflict graph. There exists a common induced incomplete poset (CIIS) P* = (*S*, ∼, ≺) *for 𝒫 if and only if S is an independent set in the conflict graph C*(𝒫).

Following Proposition 1, we can now solve the MkCIIS problem by finding a maximum independent set (MIS) in the conflict graph defined on a selection of incomplete posets. Specifically, the vertices of the independent set correspond to the subset of elements shared by at least *k* selected posets, while the independence constraint ensures that there are no conflicts for the specific selection of posets. Thus, the MIS directly yields the largest set of non-conflicting elements satisfying the requirements of the MkCIIS problem. If two elements are observed with a hidden order in some incomplete poset *P*_1_ and with an order in some incomplete poset *P*_2_, this does not cause a conflict and the two elements can be selected as part of an independent set. In the inferred recurrent trajectory, we would then use the order observed in *P*_2_ and thereby resolve the hidden order of *P*_1_. An advantage of keeping hidden orders until we find a MIS in the conflict graph is, that we do not have to determine an order before searching for recurrent trajectories. In contrast, other methods, like MASTRO [52] and RECAP [10], pull mutation clusters apart and represent the mutations in separate nodes. MASTRO connects the nodes belonging to a cluster by a special type of undirected edge. These edges are never resolved into directed edges. In RECAP, all clusters are expanded into linear paths by leveraging the information across patients and then clustered and used to build consensus trees.

To find a selection of *k* incomplete posets that yield a maximum CIIS, we propose a combinatorial optimization approach to solve the MkCIIS problem. Our algorithm consists of the following three steps:

1. We start with constructing the conflict graph for the input family of incomplete posets 𝒫 according to Definition 4.
2. We use an ILP to solve the MkCIIS problem by simultaneously selecting at least *k* incomplete posets for which an independent set in the conflict graph of the selected posets is of maximum size (Proposition 1).
3. We construct the final common trajectory by taking the incomplete subposet of the selected posets that is induced by the selected nodes.

Instead of constructing the conflict graph for each combination of incomplete posets, we construct *C*(𝒫) over the entire input and distinguish between active and non-active edges in our conflict graph when making the poset selection. An edge is active, if the two corresponding incomplete posets are selected by our model and non-active otherwise. Each edge {*i, j*} is assigned a binary variable *x*_*i,j*_ that indicates the activation of an edge. An active edge {*i, j*} prevents that *i* and *j* are both included in the node selection. Non-active edges on the contrary do not prevent the algorithm to choose both adjacent nodes.

For finding a maximum independent set in the conflict graph, here *C* = (*V*_*c*_, *E*_*c*_), we assign each node *i* a binary decision variable *x*_*i*_ that indicates whether *i* is included in the MIS or not. Similarly, each input poset *P*_*p*_ ∈ 𝒫 = {*P*_1_,…, *P*_*n*_} is represented by a binary variable *y*_*p*_ that is 1 if *P*_*p*_ is part of the solution and 0 otherwise. Further, we use a constant *a*_*i,p*_ for each node *i* in the conflict graph to denote whether *i* is contained in a poset *P*_*p*_ or not. The ILP to find maximum common trajectories in *k* posets is as follows.

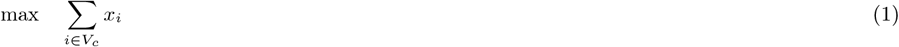

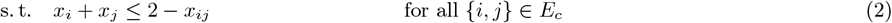

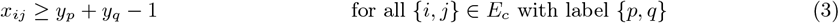

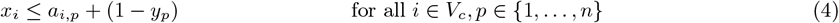

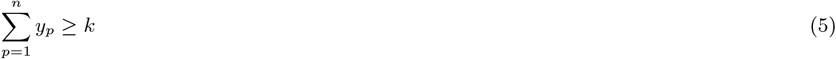

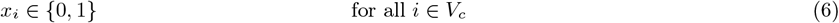

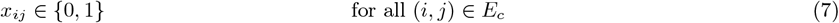

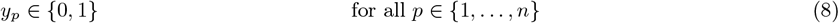

Our objective is to maximize the number of nodes that are included in the maximum independent set (1). We adapt the standard ILP formulation of the MIS problem such that no two selected nodes are connected by an active edge given a certain graph selection (2). An edge {*i, j*} with label {*p, q*} of the conflict graph is active if the corresponding posets *P*_*p*_, *P*_*q*_ ∈ 𝒫 are selected (3), thus restricting the feasible selection of nodes. Additionally, a node cannot be selected if it is not present in any selected graph (4). Finally, we require that at least *k* graphs are selected (5).

In the more general case, where the input is a family of sets of posets, we must adjust the conflict graph and the ILP formulation. The conflict graph must contain all conflicts between pairs of posets from distinct sets. In the ILP, we need an additional constraint to make a selection of at most one poset per set. We give the extended version of the conflict graph and ILP in Suppl. Mat. Sect. S1.

## 4 Results

We compare POTTR to MASTRO [52] on non-small cell lung cancer (NSCLC) [32] and acute myeloid leukemia (AML) [47] data (Section 4.1). In Section 4.2, we applied POTTR to the NSCLC TRACERx421 cohort [19], which includes phylogenetic trees inferred from multi-region whole-exome sequencing data, while Section 4.3 demonstrates the use of POTTR to identify recurrent trajectories in cell differentiation maps derived from a mouse model of development. POTTR and information on how to obtain the input data are available on GitHub https://github.com/AlBi-HHU/POTTR. We used Gurobi 12.0.0 for solving our ILP [26]. All computations were executed on a workstation with AMD EPYC 7742 64-Core processor with 128 threads, 1 TiB DDR4 RAM, running Debian Trixie with Kernel 6.12.

### 4.1 Recurrent trajectories in lung cancer and acute myeloid leukemia (AML)

We compare POTTR to MASTRO [52], since it is the only related method that evaluates recurrent trajectories as common induced subgraphs of the input, which we can compare with common induced incomplete subposets. For the comparison, we use two datasets analyzed in the MASTRO manuscript: 89 non-small cell lung cancer (NSCLC) trees inferred from multi-region whole-genome sequencing data with CITUP [46]; and 120 acute myeloid leukemia (AML) trees generated from single-cell panel sequencing data with SCITE [31] (see Suppl. Mat. Sect. S3.1 for further details). MASTRO retrieves all *maximal* recurrent trajectories that occur in at least two phylogenetic trees and applies a statistical test to evaluate the significance of each trajectory. POTTR, in contrast, identifies *maximum* recurrent trajectories in subsets of the input. For the comparison to MASTRO, we ran POTTR for every value of *k* ≥ 2 until the recurrent trajectories only included the root node, and retrieved the sizes of the maximum trajectories. We computed additional optimal solutions using Gurobi’s parameters `PoolSearchMode = 2` and `PoolSolutions = 50000`. Despite the large solution pool size, computing the maximum recurrent trajectories is relatively fast, completing in under 39 seconds for the NSCLC data and 68 seconds for the AML data for each value of *k*, whereas computing the conflicts takes about 47 seconds and 64 seconds for the NSCLC and AML data, respectively. We analyzed the runtime and memory usage on simulated data and provide the details in Suppl. Mat. Sect. S3.4.

To evaluate the significance of the trajectories reported by POTTR, we use MASTRO’s statistical test. Here, we use the null model reported in [52], that, for each tumor tree, assigns the mutations to the tree nodes independently and uniformly at random while preserving the tree topology. For each tree, the probability that a given trajectory occurs under the null hypothesis is computed individually, and the probability of observing a support that is at least as large as in the data is computed from the upper tail of the corresponding Poisson Binomial distribution [52]. Since POTTR can resolve mutation clusters, we adjusted the test to work on the trees with the resolutions found by POTTR. We refer to [52] for details of the test. Trajectories that consist of a single mutation have a *p*-value of 1.0 and are discarded by Pellegrina and Vandin [52]. Thus, in our comparison, only trajectories with at least two alterations were considered.

In the NSCLC data, POTTR and MASTRO both identify seven statistically significant maximum trajectories with *p*-values less than 0.05. POTTR detects two additional trajectories not reported by MASTRO (Fig. 4), which are obtained by resolving the order of mutation clusters. Here, we present each trajectory individually, whereas MASTRO merges them at shared nodes for visualizations. In the first trajectory, POTTR resolves the order of *COL5A2*, placing it after a cluster of mutations that includes amplification of *PIK3CA* (Fig. 4A). *PIK3CA* is a well-known cancer driver gene and a key member of the PI3K/Akt signaling pathway that regulates cell proliferation and survival. *COL5A2* is known to activate the PI3K/Akt pathway and thereby promotes resistance to erlotinib, a drug commonly used to treat NSCLC [25]. In the second trajectory (Fig. 4B), POTTR resolves the order of a cluster of mutations in *PIK3CA, TP53, SOX2*, and *NFE2L2*, inferring that mutations in *NFE2L2* occur after mutations in the other three genes. *NFE2L2* encodes the transcription factor NRF2 that plays a key role in the cellular protection against oxidative stress by regulating the expression of antioxidant proteins [27]. The increased expression of NRF2 in NSCLC has been associated with the survival of tumor cells, especially against chemotherapy [12,43]. MASTRO does identify a trajectory similar to this one, but it does not resolve the cluster, instead it reports a single cluster with a support of three containing all four mutations (Suppl. Fig. S3.2).

**Fig. 4.**
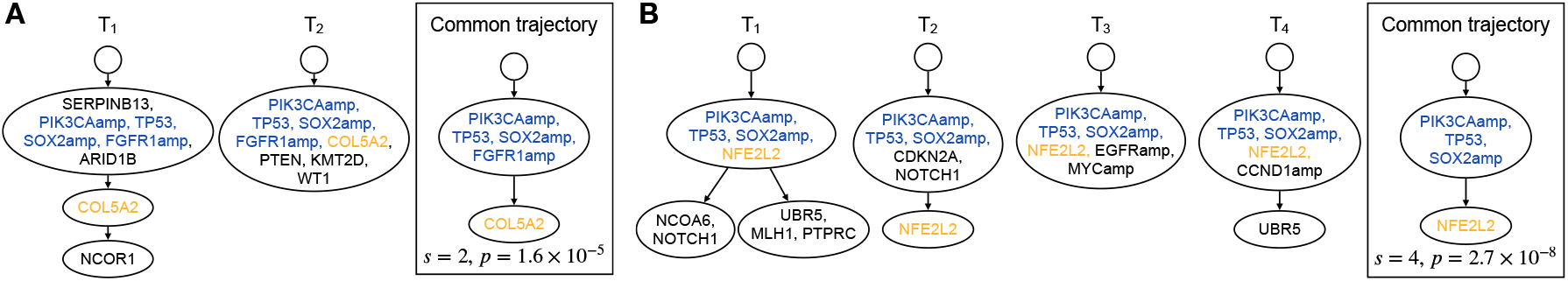
Tumor trees and corresponding recurrent trajectories (box) for maximum trajectories found by POTTR but not MASTRO in the NSCLC data with **A** support *s* = 2; **B** support *s* = 4. Corresponding *p*-values are indicated below trajectories. POTTR identified these trajectories by resolving mutation clusters based on ordering of mutations in genes *COL5A2* and *NFE2L2*. Su”x “amp” indicates amplification of corresponding gene.

POTTR and MASTRO both identify 34 maximum recurrent trajectories in the AML data, with 33 of them having a p-value below 0.05 (Suppl. Table S3.1). All of these 33 maximum recurrent trajectories identified by POTTR are statistically significant, whereas most of the maximal trajectories reported by MASTRO are not. The one trajectory with p-value > 0.05 consists of *FLT3* ≺ *NPM1*, which are the most frequent alterations in the AML dataset (Suppl. Fig. S3.1C). Overall, the recurrent trajectories found in both the AML and NSCLC datasets are small, the largest trajectories observed at *k* = 2, containing four and five alterations, respectively. As *k* increases, trajectory size decreases, reducing to single-mutation trajectories at *k* = 13 in NSCLC data and *k* = 18 in AML (Suppl. Fig. S3.1). The small trajectory sizes reflect the low per-gene mutation counts across tumors (Suppl. Fig. S3.1A and B) and low frequencies of pairwise mutation co-occurrences (Suppl. Fig. S3.3.

### 4.2 Identifying recurrent trajectories in TRACERx data

We applied POTTR to the TRACERx421 dataset, containing phylogenetic trees from 401 tumors [19] which were constructed with CONIPHER [24]. For our analysis, we only analyzed mutations with an exonic function and grouped them into two broad categories: *missense-like* (M) and *nonsense-like* (N). We classified mutations as missense-like if their exonic function was described as nonsynonymous, nonframeshift substitution, nonframeshift insertion, or splicing, while stopgain, frameshift substitution, and frameshift insertion mutations were classified as nonsense-like. We present mutations on the gene level as gene_M or gene_N in the phylogenetic trees according to the classification of the mutation. In some tumor trees, the same gene was mutated multiple times with same M, N label. Since identification of recurrent trajectories requires unique labels in trees, we retain only the first occurrence of any label. We filtered these trees to contain only mutations labeled as driver mutations in the TRACERx data, which reduced the number of distinct patients to 394 (Suppl. Mat. Sect. S3.2). CONIPHER reports multiple trees per tumor, of which some trees of an individual patient were identical after filtering. We removed duplicated trees such that each patient had a set of distinct trees. Our input consisted of 469 trees in total, with each patient contributing on average 1.2 trees that contained on average 7.7 mutations and 12.4 tree edges.

We ran POTTR on these trees, excluding MASTRO which does not have a feature to select among multiple trees from the same patient. The largest maximum trajectory contains five alterations and is supported by two trees. From *k* = 6 onward, the trajectories have only 2 mutations, and from *k* = 36 to *k* = 193, only one. For each value of *k*, POTTR identifies trajectories by resolving clusters, confirming the need of this ability. For example, we find a trajectory where *TP53* and *CDKN2A* missense mutations precede a missense mutation in *PIK3CA* (Fig. 5A), which is supported by four tumor trees. Two of them contribute to the resolution of clusters present in the other two tumor trees. The co-occurrence of *TP53* and *PIK3CA* mutations has been associated with shorter progression-free survival time [62]. Interestingly, Liao et al. [42] indicated a correlation in the TRACERx data between patients having an early *TP53* and late *PIK3CA* mutation and longer disease-free survival time compared to patients with an early *PIK3CA* mutations, that had shorter survival time and the tumors had a higher potential to spread.

**Fig. 5.**
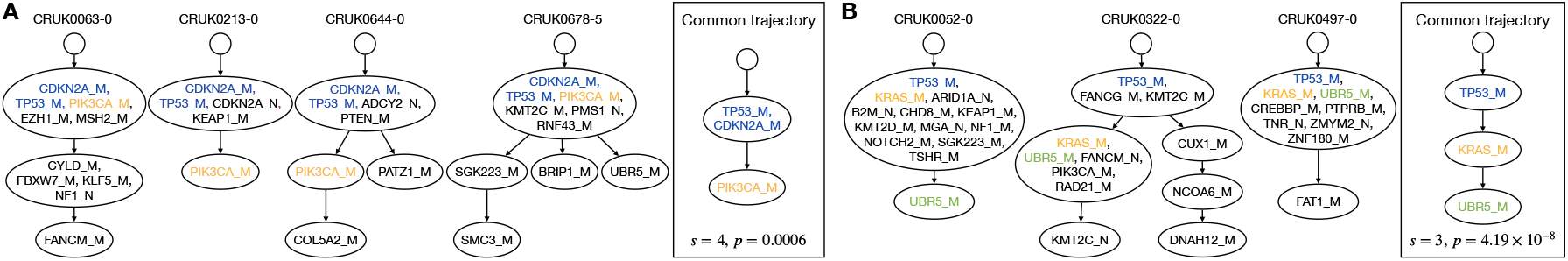
Tumor trees and corresponding maximum recurrent trajectories (box) found by POTTR through resolution of mutation clusters in the NSCLC TRACERx421 data with **A** support *s* = 4; **B** support *s* = 3. Corresponding *p*-values are indicated below trajectories. For each patient, one tree *t* was selected, indicated by −*t* in the tree label.

Another maximum trajectory containing TP53_M and KRAS_M followed by UBR5_M (Fig. 5B) illustrates POTTR’s ability to resolve multiple mutation clusters while preserving the overall partial order among mutations. The selected tree of CRUK0052 contains a cluster containing TP53_M and KRAS_M followed by UBR5_M, and in the selected tree of CRUK0322 TP53_M precedes KRAS_M and UBR5_M (Fig. 5B). These clusters are resolved such that the inferred order TP53_M→ KRAS_M→ UBR5_M is maintained.

### 4.3 Recurrent cell differentiation trajectories in Trunk-Like Structures (TLS) – an in vitro model of the mammalian embryonic trunk

We use POTTR to identify conserved cell differentiation routes during mammalian trunk development and how these routes change due to chemical perturbations. Specifically, we analyzed cell differentiation maps derived from single-cell CRISPR-Cas9-based lineage tracing of an *in vitro* embryoid model called Trunk-like Structures (TLS) [5]. We used POTTR to compare unperturbed TLS to TLSCL, a perturbed system treated with a WNT activator (CHIR99021) and a BMP inhibitor (LDN193189) during the developmental process.

A cell differentiation map is a directed acyclic graph (DAG) whose vertices represent cell types and whose edges represent transitions (differentiation events) between cell types that occur during development. Specifically, vertices with no outgoing edges, i.e. sinks of the cell differentiation map, represent terminal cell types, while the internal vertices with outgoing edges represent progenitor cell types. Each progenitor cell type is represented by its potency, or the set of terminal cell types that can be attained by the descendants of the progenitor. The dataset comprises 12 TLS maps and 11 TLSCL maps, each corresponding to an independent biological replicate and containing six cell types – endoderm, endothelial, NMPs, neural tube, PCGLC, and somite. Differentiation maps were constructed using CARTA [56] for each biological replicate independently (see Suppl. Mat. Sect. S3.3 for details). Since cell differentiation maps are, in general, DAGs rather than trees, existing methods designed to analyze trees are not equipped to detect recurrent developmental trajectories. POTTR’s ability to compare and extract shared patterns directly from general directed acyclic graphs makes it uniquely suited for this analysis.

POTTR identifies TLS differentiation routes that are conserved across replicates and how these routes are altered under WNT activation and BMP inhibition. We examined the differentiation maps inferred by POTTR that are conserved in more than half of the TLS and TLSCL biological replicates, respectively. While the differentiation maps, containing six and five cell types, are broadly similar, they diverge in the inferred origin of somites (Fig. 6). In the canonical view of trunk development, somites arise from progenitors that share ancestry with neural tube cells. However, the conserved TLS differentiation map highlights an alternative trajectory in which somites emerge from a progenitor that also gives rise to endothelial cells, shown in yellow in Fig. 6 A. This is consistent with previous *in vivo* studies that have found evidence for a secondary pathway towards the production of the trunk endothelium [40,48]. In contrast, TLSCL differentiation map infers the stereotypical route in which somitic cells originate from neuro-mesodermal progenitors, indicating that WNT activation and BMP inhibition suppresses the alternative somite–endothelium route of somitic differentiation.

**Fig. 6.**
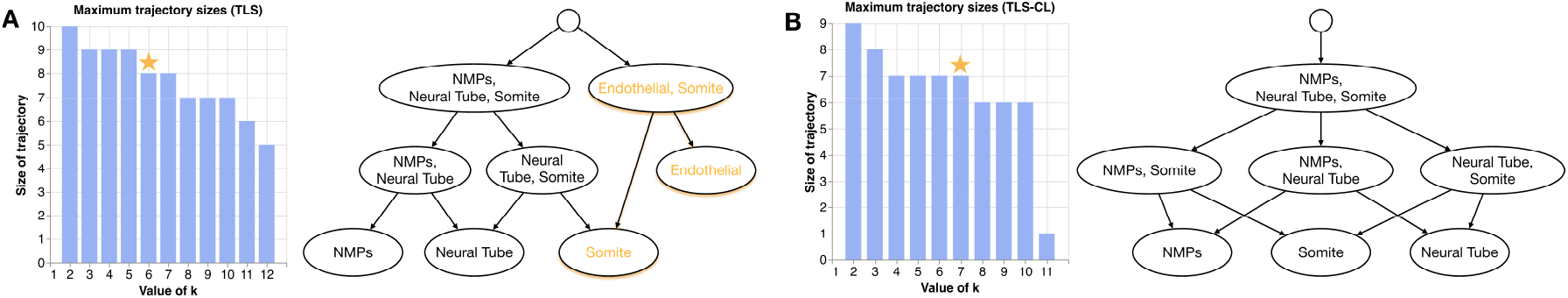
Size histogram of maximum recurrent trajectories in mouse cell differentiation maps for different values of the parameter *k* and maximum recurrent trajectories found by POTTR for *k* = 6 and *k* = 7 (star) in **A** TLS and **B** TLSCL differentiation routes.The alternate route through which somites emerge is highlighted in the TLS conserved differentiation map.

## 5 Discussion

We formulated the problem of finding a recurrent trajectory as the problem of finding a common induced incomplete subposet, and derived an algorithm POTTR to find such posets even in cases where one must select among multiple possible orders in each sample. This rigorous formulation generalizes to arbitrary partial orders (not only trees), naturally describes the problem of resolving orders of events that are indistinguishable in some samples, and avoids heuristics involving tree operations that are used in many existing recurrent trajectory methods. We demonstrated these advantages on two cancer sequencing datasets and one lineage tracing dataset from mouse development. POTTR scales to large datasets including the TRACERx data [32,19] containing 469 trees from 394 patients. In contrast, MASTRO enumerates all maximal trajectories, which becomes intractable for a large input with many conserved trajectories. POTTR also resolves mutation clusters which is essential when analyzing bulk-sequencing data where there is often considerable uncertainty in the trees.

There are a number of limitations of the current approach and analyses. First, in current datasets the support for resolving mutation clusters is small (1-2 patients) and might be due to errors in tree construction. Larger sample sizes, phylogenetic trees with a higher accuracy, or a combination of bulk and single-cell sequencing data might improve the confidence when resolving clusters. At the same time, finding common induced incomplete subposets is an NP-hard problem and thus finding recurrent trajectories from larger datasets might require better heuristics and approximation algorithms, possibly by adapting algorithms for the maximum independent set problem [6,17,9]. Second, assessing the statistical significance of recurrent trajectories remains challenging. We are currently using MASTRO’s permutation test [52], but have observed that the maximum recurring trajectories reported by POTTR were almost always statistically significant. Further, this significance test is only feasible if the number of nodes is lower than approximately ten, since the test analyzes all automorphisms of a trajectory. Extending the poset approach to a probabilistic setting [44,45] is an interesting future direction. Third, similar to most methods, we apply the infinite sites assumption, stating that each mutation is gained once and from then on inherited to all descendants. This enables a clear identification of common mutation events. However, it is not unusual to observe parallel mutation processes in distinct subclones [47]. While lifting the infinite sites assumption is more realistic, it complicates identifying recurrent trajectories, as suddenly a mutation event might have multiple corresponding events across the data. To partially overcome this assumption, we have introduced a new labeling system that allows multiple mutation events per gene in the incomplete posets. Removing the infinite sites assumption completely is left for future work.

Finally, there are some unexplored connections between the problem of finding recurrent trajectories and computing tree edit distance [59]. Several well-studied problems for comparing trees can be viewed as restricted or special cases of the tree edit distance problem [4,33,38,35]. Finding a maximum common trajectory between two vertex-labeled trees is yet another restricted form of the tree edit distance problem, one in which all deletions must precede any insertions, and no substitutions are allowed. Further investigation of these connections might yield better algorithms for specialized tree edit distance or recurrent trajectory problems.

