## Supplementary Material for "POTTR: Identifying Recurrent Trajectories in Evolutionary and Developmental Processes using Posets"

### Supplement S1. Finding recurrent trajectories in a family of sets of posets

Reconstructing the phylogeny of a tumor from bulk sequencing data is challenging due to the tumors inherent heterogeneity and limitations of bulk sequencing technologies, which lead to many plausible phylogenies. Methods that compute phylogenetic trees from bulk sequencing data often report multiple trees per tumor [1]. In a more realistic setting, the input to POTTR consists of a family of phylogenetic trees for each tumor. However, a tumor arises from one clonal evolutionary history initiated by a single cell-of-origin that undergoes genetic alteration. Thus, we must decide on one phylogeny per tumor, which we can use to identify recurrent evolutionary trajectories.

Considering the ambiguity of the data, we focus on finding a maximum common induced subposet among $k$ selected sets from the family $\mathscr{P}$, where $k$ is a user-defined integer. From each selected set, one poset is chosen such that the common induced subposet is an induced subposet of either the chosen poset or one of its resolutions. We formally pose this generalized version of the MkCIIS problem as follows.

**Problem 2 (Maximum $k$-Common Induced Incomplete Subposet Problem, (MkCIIS))** *Given a family $\mathscr{P} = \{\mathcal{P}_1, \ldots, \mathcal{P}_n\}$, where each $\mathcal{P}_i$ is a set of posets, and an integer $k$, find a largest poset $P$ and a selection of at least $k$ sets of $\mathscr{P}$, with one poset selected from each, such that $P$ is an induced subposet of the selected poset or one of its resolutions.*

We adjust the conflict graph $C(\mathscr{P})$ such that it encodes conflicts in the order of events described by any pair of posets from distinct sets in $\mathscr{P}$. We formally define the conflict graph $C(\mathscr{P})$ for $\mathscr{P}$ as follows.

**Definition 5 (Conflict Graph).** *A conflict graph $C(\mathscr{P})$ of a family $\mathscr{P} = \{\mathcal{P}_1, \ldots, \mathcal{P}_n\}$, where each $\mathcal{P}_i = \{(S_i, \sim_i^{(1)}, \prec_i^{(1)}), \ldots, (S_i, \sim_i^{(m_i)}, \prec_i^{(m_i)})\}$ is a set of posets, is an undirected multigraph. The vertex set consists of all elements that occur in at least two distinct sets: $V = \{v \in \bigcup_{i=1}^{n} S_i \mid |\{i \mid v \in S_i\}| \geq 2\}$. There is an edge $\{a, b\}$ with label $i_1, i_2$ in $C(\mathscr{P})$ for each pair of posets $(S_{i_1}, \sim_{i_1}^{(j_1)}, \prec_{i_1}^{(j_1)})$ and $(S_{i_2}, \sim_{i_2}^{(j_2)}, \prec_{i_2}^{(j_2)})$ from distinct sets $\mathcal{P}_i$ and $\mathcal{P}_j$ if any of the following conflicts exists: (i) $a \prec_{i_1}^{(j_1)} b$ and $b \prec_{i_2}^{(j_2)} a$; or (ii) $a \prec_{i_1}^{(j_1)} b$ and $a \parallel_{i_2}^{(j_2)} b$; or (iii) $a \sim_{i_1}^{(j_1)} b$ and $a \parallel_{i_2}^{(j_2)} b$.*

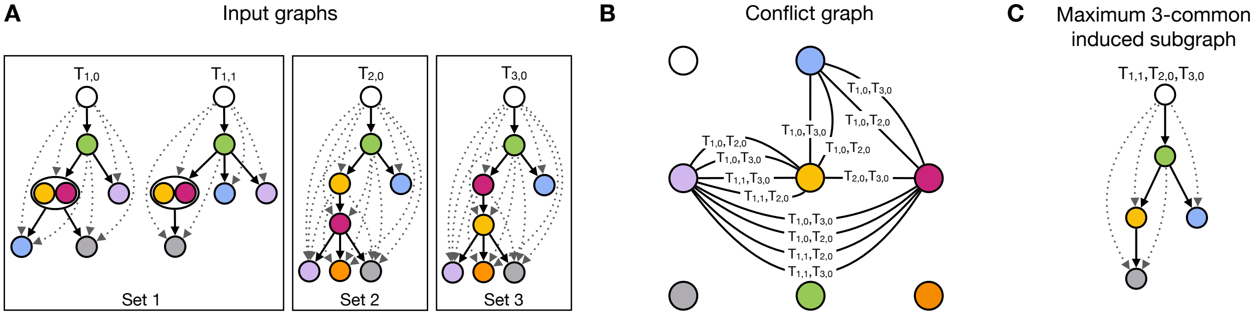

**Fig. S1.1.** Example of the conflict graph for three sets of input posets. **A** Three distinct sets of input posets presented as transitively closed DAGs. Set 1 consists of two posets, which both have a hidden order for the yellow and pink element. The hidden orders are represented as clusters in the DAGs. **B** Conflict graph with all pairwise conflicts between posets of distinct sets. **C** A maximum common induced subposet for the selection of $T_{1,1}, T_{2,0}$, and $T_{3,0}$.

Figure S1.1 gives an example of the conflict graph of a family of poset sets and a maximum common induced subgraph. For visualization, again, we present posets as transitively closed DAGs in panel A. We give a maximum common induced subgraph in Figure S1.1C for $k = 3$ selected input sets, for which one poset is selected.

We incorporate this decision procedure into our ILP formulation, obliquely making the selection of phylogenies part of the optimization. Consequently, the choice of phylogenetic trees can affect the size of a maximum independent set in the conflict graph, and thereby determine the size of the resulting recurrent trajectory.

We compute all pairwise conflicts from all posets in one set to all other posets of the other sets. The ILP formulation is equal to the previous one, where the input consisted of one poset per evolutionary process, with some minor changes. Just as before, all conflicts are added to the conflict graph $C = (V_c, E_c)$ with labels indicating the conflicting poset pairs. Each phylogeny $t$ related to a patient $p$ is represented by a binary variable $y_{pt}$. In our updated ILP formulation below, we account for multiple trees per evolutionary process, e.g., tumors, and add a constraint to select at most one phylogeny per tumor.

$$
\begin{aligned}
\max \quad & \sum_{i \in V_c} x_i && && \text{(S1.1)} \\
\text{s. t.} \quad & x_i + x_j \leq 2 - x_{ij} && \text{for all } (i,j) \in E_c && \text{(S1.2)} \\
& x_{ij} \geq y_{p_1 t_1} + y_{p_2 t_2} - 1 && \text{for all } (i,j) \in E_c : && \text{(S1.3)} \\
& && (i,j) \in c((p_1,t_1),(p_2,t_2)) && \\
& x_i \leq a_{i,pt} + (1 - y_{pt}) && \text{for all } i \in V_c, p \in P, t \in T_p && \text{(S1.4)} \\
& \sum_{t \in T_p} y_{pt} \leq 1 && \text{for all } p \in P && \text{(S1.5)} \\
& \sum_{p \in P} \sum_{t \in T_p} y_{pt} \geq k && && \text{(S1.6)} \\
& x_i \in \{0,1\} && \text{for all } i \in V_c && \text{(S1.7)} \\
& x_{ij} \in \{0,1\} && \text{for all } (i,j) \in E_c && \text{(S1.8)} \\
& y_{pt} \in \{0,1\} && \text{for all } p \in P, t \in T_p && \text{(S1.9)}
\end{aligned}
$$

Constraints S1.2-S1.4 are equivalent to the constraints before. Since we have multiple input posets per evolutionary process, we must make sure to select at most one per evolution S1.5. When selecting the $k$ input posets, we must also account for having multiple posets per process now S1.6.

### Supplement S2. Proofs

#### S2.1 Proving the NP-hardness of the MkCIIS problem

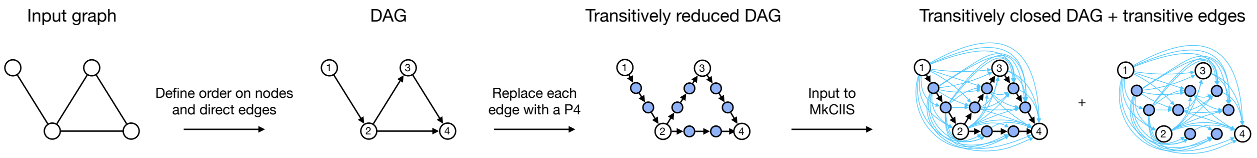

**Fig. S2.1.** Steps to transform the input graph of the maximum independent set problem into an input instance of the maximum $k$-common induced incomplete subposet problem. The transformation starts with an undirected graph and results in two transitively closed DAGs that are equivalent to posets.

To show that the decision variant of the MkCIIS problem is NP-complete even for $k = 2$, we use a reduction from the maximum independent set (MIS) problem. To transform an instance of the MIS problem to an instance of the MkCIIS problem, we apply several steps, shown in Figure S2.1.

*Proof.* The decision variant of the MkCIIS problem has an additional parameter $s$ and asks, if a maximum common induced incomplete subposet of size at least $s$ exists for a selection of at least $k$ incomplete posets.

For a given solution $P = (S, \prec, \sim)$ and a set of selected incomplete posets $\mathscr{P}$, we can verify in polynomial time if $\mathscr{P}$ has at least $k$ elements and if $|S| \geq s$. Further, we must verify that $P$ is a common induced incomplete subposet, which we do in polynomial time as follows. We create the induced incomplete subposet on $S$ of each selected incomplete poset and compare it to $P$. Since we must compare all pairwise relation, we check $|S|^2$ relations for at least $k$ selected posets, taking $\mathcal{O}(k|S|^2)$ time. Hence, the problem is in NP.

We now give a reduction of the well-known NP-hard MIS problem [21] Its decision variant asks if for a given undirected graph $G = (V, E)$ and a parameter $s'$, is there an independent set $V' \subseteq V$ of size $s'$? Given such an instance of the MIS problem, we transform it to an instance of the MkCIIS problem described as follows and depicted in Figure S2.1. As the MkCIIS problem is defined on transitively closed DAGs, we start with defining an increasing order on $V$ and make $G$ a DAG by directing the edges such that small nodes have outgoing edges towards larger nodes. Next, we use the Poljak Lemma [53], where each edge in $G$ is replaced by a path of three edges, i.e., a $P_4$, see panel with blue nodes in Figure S2.1. The resulting graph $F$ is transitively reduced and the independence number of $F$ is increased by the number of edges in the original $G$, $\alpha(F) = \alpha(G) + |E|$ [53]. This is, because we can extend each independent set of $G$ in $F$ by one node from each added $P_4$. Finally, to get the input of the MkCIIS problem, we make $F$ transitively closed and define a second graph $H$ on the same node set $V$ with all the transitive edges, that we added to $F$. Due to the one to one correspondence of transitively closed DAGs and posets, we can use $F$ and $H$ as input to our MkCIIS problem, where we represent them as incomplete posets, where the hidden order is empty. We can duplicate $H$ multiple times to show hardness for any value of $k \geq 2$. For now, we take $F$ and $H$ once, and set $k = 2$ and $s = s' + |E|$. All transformations of the input of the MIS problem into the input of the MkCIIS problem take polynomial time.

By definition of the MkCIIS problem, here, we search for a set of elements with the same pairwise orders in $F$ and $H$. In terms of graph language, we search for a graph that is an induced subgraph in the two graphs. Since $H$ does not contain any of the non-transitive edges of $F$, the black edges in the last panel of Figure S2.1, the CIIS does not contain any non-transitive edges either. Any solution of the MkCIIS problem contains exactly one of the additional nodes in $F$ for each edge replaced by a $P_4$. If the MIS problem has a solution of size $s'$, then the MkCIIS problem has a solution of size $s' + |E|$ for $k = 2$. Vice versa, if we have a solution of size $s' + |E|$ for the MkCIIS problem, when removing all transitive edges and the $|E|$-many additionally inserted nodes, the result is an independent set in $G$ of size $s'$ and thereby a solution of the MIS problem. It follows, that the MIS instance has a yes-answer if and only if the transformed MkCIIS instance has a yes-answer too. □

#### S2.2 Independent sets in the conflict graph

Here we give proof of Proposition 1, which gives the correspondence of independent sets in the conflict graph and recurrent trajectories in the input incomplete posets.

*Proof.* The orders and incomparabilities of a CIIS $P = (S, \sim, \prec)$ of a family $\mathscr{P}$ of incomplete posets are, by definition, shared by all incomplete posets in $\mathscr{P}$. Consequently, the elements of $S$ have no conflicting relations and thus no edges in the conflict graph $C(\mathscr{P})$, meaning that $S$ forms an independent set in $C(\mathscr{P})$.

Edges in the conflict graph $C(\mathscr{P}) = (V, E)$ indicate conflicting relations. By definition, none of the vertices of an independent set are adjacent in a graph. Therefore, any independent set $V' \subseteq V$ in $C(\mathscr{P})$ is a set of elements without conflicts and thereby defines a CIIS $P = (V', \prec, \sim)$ of $\mathscr{P}$ on the vertex set $V'$. □

### Supplement S3. Additional experimental results

#### S3.1 Comparison MASTRO

We compared POTTR with MASTRO on NSCLC and AML datasets provided by [52]. Originally, phylogenetic trees for 99 NSCLC patients [32] and 123 AML patients [47] are available, with multiple plausible phylogenies for some patients. The MASTRO paper chose one tree per patient and removed trees with fewer than two alterations. Mutations are presented on gene-level in the trees, except for gene amplifications, which Pellegrina and Vandin kept as a distinct alteration type [52].

#### S3.2 TRACERx experiments

When we filtered for mutations labeled as driver mutations in the TRACERx data, of the 401 trees only 394 remained. We observed that the patients CRUK0193, CRUK0715, CRUK0118, CRUK0764, CRUK0343 had no mutations labeled as driver mutations. CRUK0586_Tumour1 and CRUK0386 each had one driver mutation, however, they were not assigned in their trees. Therefore, we removed these trees from our first analysis.

**Table S3.1.** Overview of maximum trajectories found by POTTR and MASTRO in the NSCLC and AML data. For each number of trees sharing a recurrent trajectory, only the trajectories of maximum size were considered.

| | NSCLC | | | AML | | |
|---|---|---|---|---|---|---|
| | sign. $\leq 0.02$ | sign. $\leq 0.05$ | all | sign. $\leq 0.005$ | sign. $\leq 0.05$ | all |
| Shared | 6 | 7 | 7 | 27 | 33 | 34 |
| Unique to POTTR | 2 | 2 | 2 | 0 | 0 | 0 |
| Unique to MASTRO | 0 | 0 | 0 | 0 | 0 | 0 |

**Table S3.2.** Overview of trajectories found by POTTR and MASTRO in the NSCLC and AML data. POTTR reports trajectories of maximum size, while MASTRO reports all maximal trajectories.

| | NSCLC | | | AML | | |
|---|---|---|---|---|---|---|
| | sign. $\leq 0.02$ | sign. $\leq 0.05$ | all | sign. $\leq 0.005$ | sign. $\leq 0.05$ | all |
| Shared | 6 | 7 | 7 | 27 | 33 | 34 |
| Unique to POTTR | 2 | 2 | 2 | 0 | 0 | 0 |
| Unique to MASTRO | 9 | 18 | 116 | 13 | 36 | 103 |

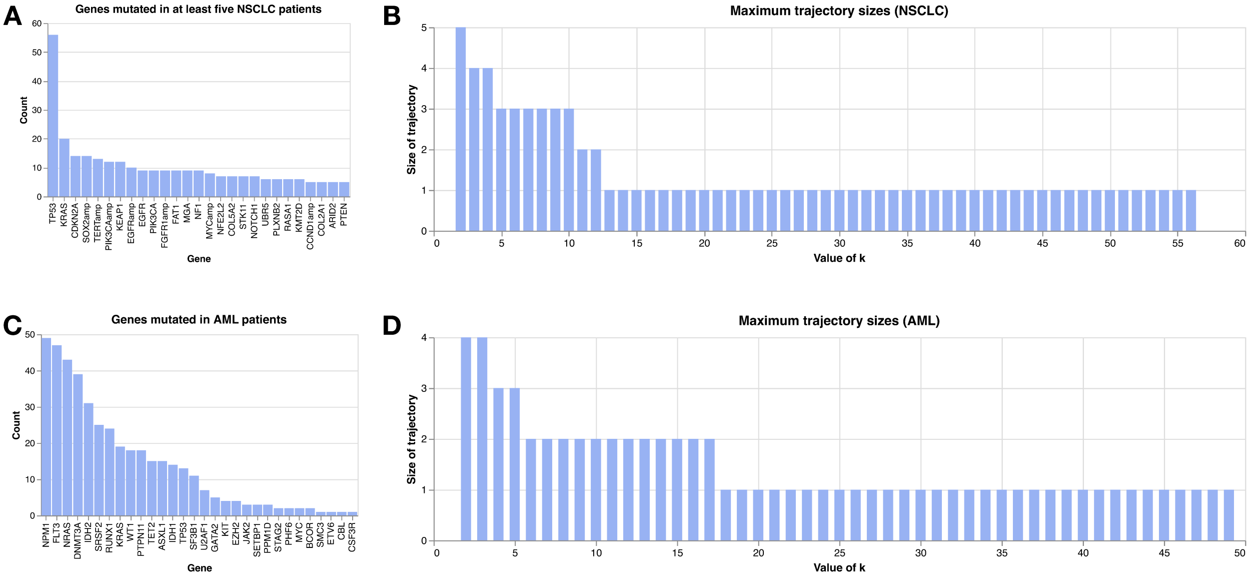

**Fig. S3.1. A** Most common mutations in NSCLC data. Due to high mutation diversity, we only display genes that are mutated in at least five distinct patients. **B** Number of mutations in the maximum trajectories for every $2 \leq k \leq 56$, for any larger $k$ the trajectory only included the root node. **C** Most common mutations in AML data. **D** Number of mutations in the maximum trajectories for every $2 \leq k \leq 49$, for any larger $k$ the trajectory only included the root node.

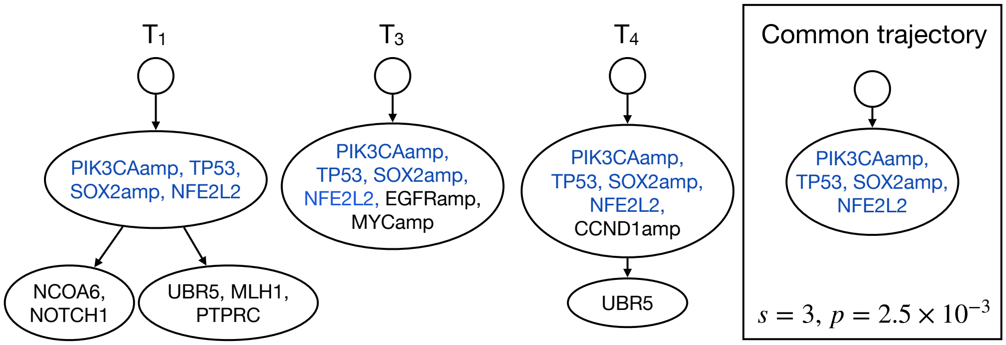

**Fig. S3.2.** Three graphs for which MASTRO identifies a common trajectory, presented in the box with the support and p-value, that consists of a single mutation cluster.

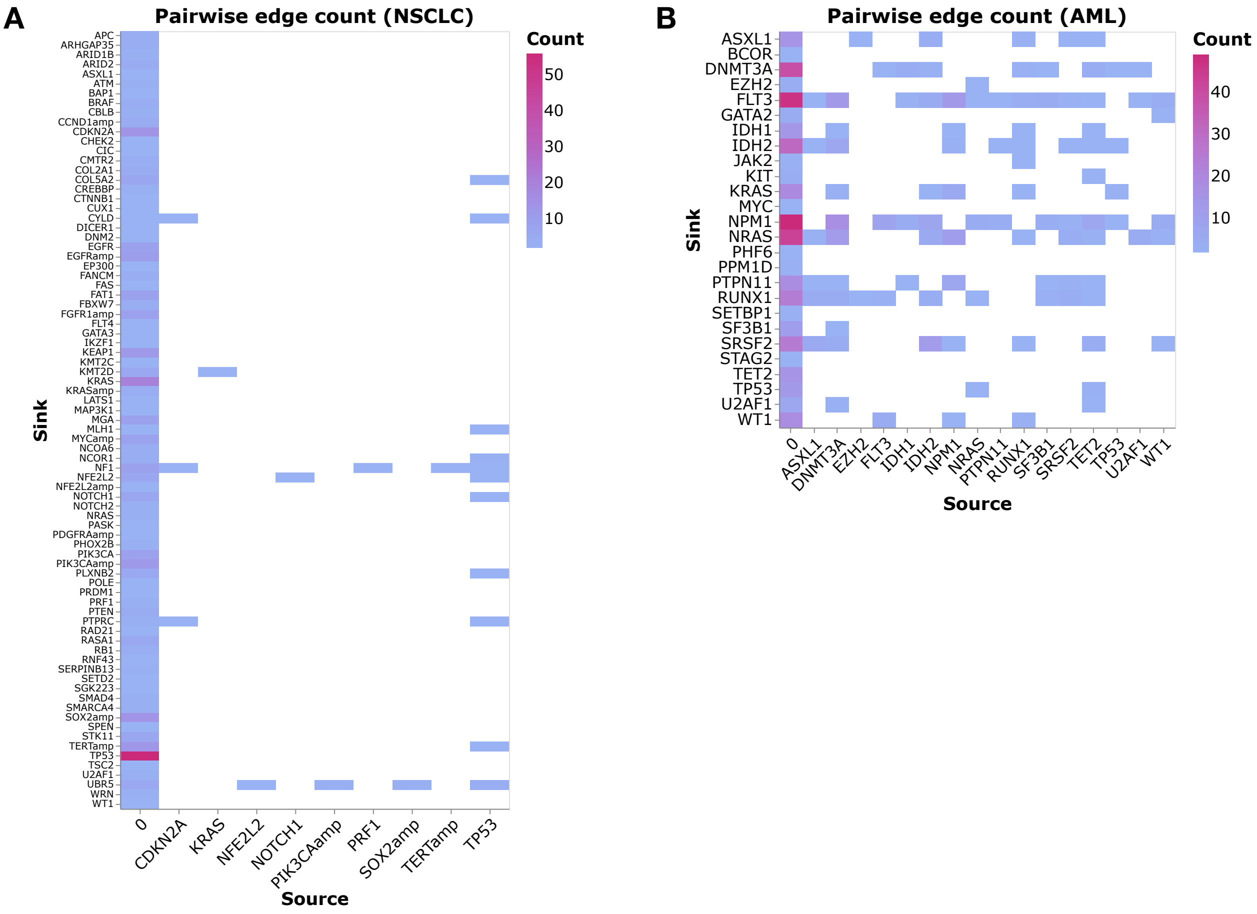

**Fig. S3.3.** Absolute count of edges that occur at least in two graphs in the **A** NSCLC data with a maximum of 56 and **B** AML data with a maximum of 49. Edges have the direction (Source, Sink). The source 0 stands for the root node. Mutations that are co-occurring in a cluster were not considered here.

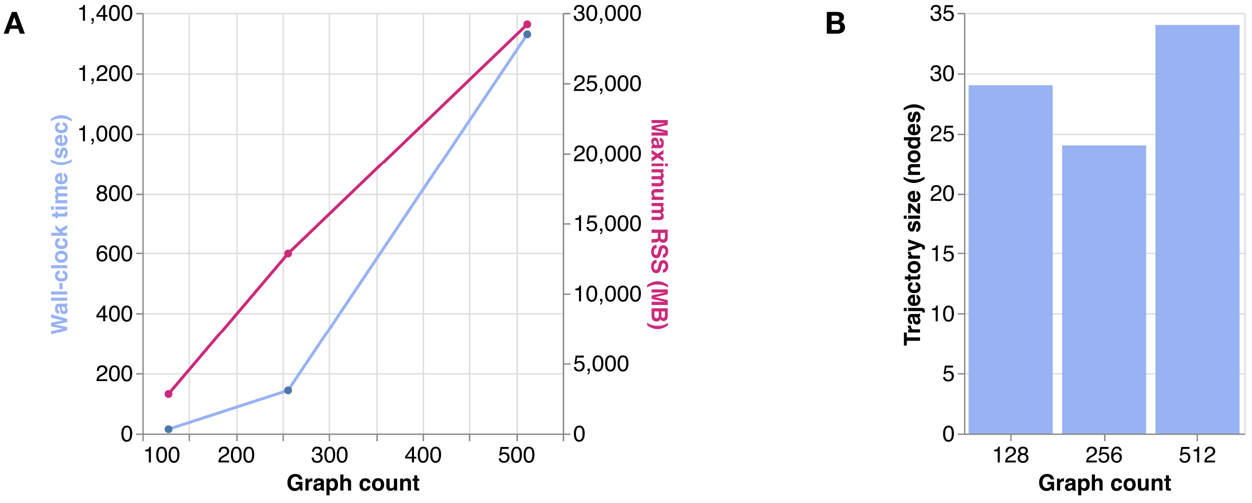

**Fig. S3.4.** Benchmark results for three instances of simulated data. **A** Wall-clock time in seconds (blue) and maximum resident set size in megabytes (pink). **B** The sizes of the trajectories found by POTTR for the three instances.

#### S3.3 Cell differentiation experiments

Cell differentiation maps by labeling each internal vertex of the cell lineage trees by its observed potency, i.e. the set of cell types observed in the subtree rooted at the vertex. CARTA [56] uses these observed potencies to create cell differentiation maps where nodes represent progenitor or terminal cell types and edges represent cell type transitions. Further, CARTA annotates edges by the number of cells that follow this cell type transition [56]. To increase the confidence in the cell differentiation maps, we filtered out edges with a weight of one unless it would make the DAG unconnected.

#### S3.4 Benchmark results

We simulated tumor trees using the observed edge frequencies of the TRACERx data set described in Section 4.2. We individually created instances with 128, 256, 512, and 1024 graphs with on average 19.73 nodes and 34.73 edges including the edges added through transitive closure. The minimum and maximum number of edges were one and 161 (for the set of 512 graphs), respectively. The simulated data is thereby larger than all real data sets under consideration. We measured the runtime and peak amount of RAM (maximum resident set size (RSS)) used by POTTR (Suppl. Fig. S3.4). For our benchmarking, we did not execute the statistical significance test by MASTRO. The benchmarks were performed on our workstation using 128 threads and a time limit of ten hours. Within the time limit, the instance with 1024 graphs did not finish. The trajectories reported in Suppl. Fig. S3.4B were computed for $k = 2$.

### Supplement S4. Notation overview

| **Notation** | **Explanation** |
|---|---|
| $\prec$ | partial order, which is reflexive, asymmetric, and transitive |
| $a \prec b$ | a precedes b |
| $P = (S, \prec)$ | partially ordered set (poset) |
| $a \sim b$ | hidden order between a and b, i.e., a precedes b **or** b precedes a, but not both |
| $(\sim, \prec)$ | incomplete partial order |
| $P = (S, \sim, \prec)$ | incomplete partially ordered set (poset) |
| $P_{\mathcal{H}}$ | partition induced by $\sim$ on $S$, the sets of the partition are related by a strict partial order |
| $\mathscr{P}$ | family of posets |
| $C(\mathscr{P})$ | conflict graph defined on the family of posets $\mathscr{P}$ |
